# Thinner is not always better: Optimising cryo lamellae for subtomogram averaging

**DOI:** 10.1101/2023.07.31.551274

**Authors:** Maarten W. Tuijtel, Jan Philipp Kreysing, Sonja Welsch, Gerhard Hummer, Martin Beck, Beata Turoňová

## Abstract

Cryo-electron tomography (cryo-ET) is a powerful method to elucidate subcellular architecture and to structurally analyse biomolecules *in situ* by subtomogram averaging (STA). Specimen thickness is a key factor affecting cryo-ET data quality. Cells that are too thick for transmission imaging can be thinned by cryo-focused-ion-beam (cryo-FIB) milling. However, optimal specimen thickness for cryo-ET on lamellae has not been systematically investigated. Furthermore, the ions used to ablate material can cause damage in the lamellae, thereby reducing STA resolution. Here, we systematically benchmark the resolution depending on lamella thickness and the depth of the particles within the sample. Up to ca. 180 nm, lamella thickness does not negatively impact resolution. This shows that there is no need to generate very thin lamellae and thickness can be chosen such that it captures major cellular features. Furthermore, we show that gallium-ion-induced damage extends to depths of up to 30 nm from either lamella surface.

## Introduction

Cryo-electron tomography (cryo-ET) enables 3-dimensional observation of near-natively preserved cryogenically fixed biological specimens^1,2^. When followed by subtomogram averaging (STA)^3,4^, *in situ* cryo-ET combines high-resolution structural information of proteins or macromolecular complexes with contextual information within their functional crowded cellular environment^5,6^. Apart from technical specifications of the imaging equipment, the quality of cryo-ET datasets is greatly influenced by the properties and quality of the sample. Due to the limited penetration power of electrons, sample thickness is a particularly major limitation^7^. The influence of sample thickness on resolution is well established for single-particle cryo-electron microscopy (SPA cryo-EM)^8,9^. Although many biological samples are thin enough to be imaged directly^10–12^, other samples, most notably eukaryotic cells, need to be thinned prior to imaging^13^. Currently, cryo-focused-ion-beam milling (cryo-FIB milling) is the thinning method of choice, as it lacks several severe artefacts associated with cryo-sectioning^14,15^. During cryo-FIB milling, material from the sample is ablated by focused ions, until a thin (up to ca. 300 nm) slice remains, which can subsequently be subjected to tomographic acquisition^16^. The cryo-FIB to STA workflow has been successfully applied to a wide variety of samples and recently high-resolution maps have been obtained with local resolutions up to 2.4 – 3.5 Å^17–19^. It is known that samples amass structural damage along their milling surface during cryo-FIB milling, induced by implanted ions or cascading secondary particles resulting from interactions with ions^15,20^. In material sciences, this damage can be directly observed and has been quantified for many materials, ionic species and ion energies^21–24^. These results have previously been extrapolated to biological material^15^, recently experimentally tested for argon^20^ and gallium^25^ ions. The latter study used a 2D template matching approach^26^ to assess damage, however notably no STA or averaging approaches were used to quantify structural damage from cryo-FIB milling. While the effect of cryo-ET dose-distribution on the resolution attained by STA has previously been benchmarked^27^, the effect of the use of lamellae and their properties on cryo-ET and STA has not yet been systematically investigated. Here we benchmark the effect of the local lamella thickness on STA-resolution, and quantify the extent and degree of structural damage caused by gallium ions during the cryo-FIB milling process. By milling thinner lamellae, the contrast and signal-to-noise-ratio in the tilt-images increases, likely resulting in better alignment of the tilt-series prior to tomogram reconstruction, which is a crucial step that determines the quality of the tomogram and thereby all successive processing steps. Simultaneously however, very thin lamellae are non-trivial to prepare, contain less of the cellular context, and larger particles may no longer completely fit inside, hindering their identification. Moreover, the volume that remains undamaged by ions is concomitantly smaller as well, resulting in a trade-off between contrast and signal-to-noise-ratio on one side and the ion-damage layer on the other. Here, we used ribosomes as a probe to determine the influence of lamella thickness on STA resolution and to assess the effect of ion-induced-damage by gallium ions during cryo-FIB milling. We found that intermediate lamellae thickness up to 180 nm and particles that are located at least 30 nm into the lamellae can be used without any loss of resolution. Overall, our systematic benchmark provides guidance for the milling of lamellae and the selection of particles to optimise resolution for STA.

## Results

For this study, we extended a previously collected cryo-ET dataset of *Dictyostelium discoideum* cells^17^ to a total of 261 tilt-series collected from 13 lamellae, by acquiring additional tilt-series on a Titan Krios G4 equipped with a cold field emission gun, Selectris X energy filter (set to a slit width of 10 eV) and a Falcon 4 camera, at a pixel size of 1.2 Å (also see “Methods”). After pre-processing and tomogram reconstruction in IMOD^28^ (see “Methods”), the thickness of the lamella within each tomogram was measured manually, which will be referred to as the local lamella thickness. We found that local lamella thickness ranged from 43 – 255 nm (see Fig. 1a). Due to the geometry of the ion beam during the cryo-FIB milling process, local lamella thickness increases from the front towards the back of the lamella, creating a wedge-shape (see Fig. 1b,c)^29^. Tomograms were reconstructed again using Warp^30^ and ribosome particles were identified by template matching in STOPGAP^31^. We extracted ca. 168,000 particles as putative ribosomes, which are hereafter referred to as the raw template matches. To filter out false positives, these particles were subjected to multiple rounds of 3D classification on binned data in Relion 3.1^32^, after which ca 48,000 ribosomal particles were retained for further analysis, see Supplementary Fig. 1.

**Fig. 1.**
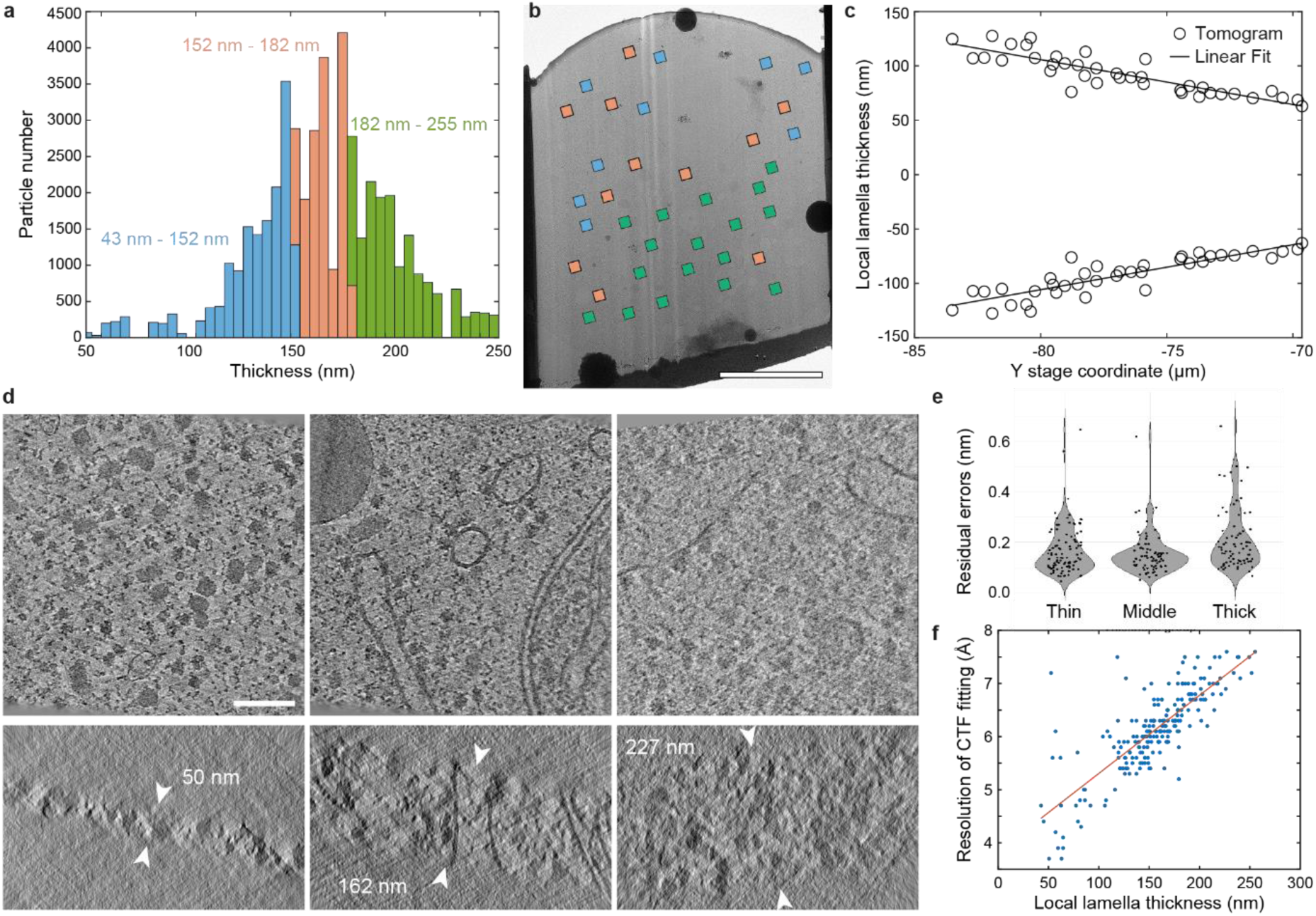
Cryo-tomography dataset overview. **a** Histogram of the local lamella thickness based on all subtomograms in the dataset. These were divided into three thickness groups, each containing 16,114 subtomograms. **b** Typical lamella imaged in a cryo-transmission electron microscope. Squares represent position and field-of-view of acquired tilt-series, coloured according to the local thickness scale shown in a. **c** Projected local lamella thickness from front (right) to back (left) of the lamella shows the wedge shape due to cryo-FIB geometry. Local thickness was divided by 2, and mirrored towards negative Y-scale for visualisation. **d** XY (top) and XZ slices (bottom) through reconstructed tomograms from each thickness group, displaying local thickness of lamella between the white arrowheads. **e** Violin plot of the residual errors after tilt-series alignment for each thickness group. **f** Resolution up to which CTF was reliably fitted in Warp for every tilt-series in the dataset plotted against the local thickness of the lamella. The red line represents a linear fit through all data points. Scalebars: 5 µm in b, and 100 nm in d (applies to all subpanels).

### Averaging ribosomes from lamellae with local thickness up to ca. 180 nm has no adverse effect on resolution

To study the effect of local lamella thickness on the resolution obtained by STA, we divided all subtomograms of ribosomes in equally sized groups, based on the local lamellae thickness. This resulted in the three thickness groups: 43 – 152 nm (“thin”), 152 – 182 nm (“middle”) and 182 – 255 nm (“thick”) (See Fig. 1a, d), each containing 16,114 particles. Beyond cryo-FIB damage and reduced electron penetration power due to increased sample thickness, tilt-series alignment is of critical importance to the quality of the tomograms^33^. Cryo-FIB-milled lamellae lack high-contrast fiducial markers and the contrast is further reduced by the crowded cellular interior, which is exacerbated for thicker lamellae. Surprisingly, we find that the residual errors after tilt-series alignment only vary slightly for the three thickness groups (See Fig. 1e). Another important parameter to consider is the fitting of the contrast-transfer function on the tilt-images; the precision of which is dependent on the thickness of the sample. Warp estimates the resolution up to which the CTF can be reliably fitted^30^, and we found a linear dependency on this estimated resolution against the local thickness, see Fig. 1f.

All ribosomal particles were extracted using Warp^30^ and subjected to 3D refinements in Relion 3.1^32^. To quantify the quality of data coming from each of the thickness groups, we analysed subsets of each thickness group to plot square inverse resolution versus particle numbers (a so-called Rosenthal & Henderson plot^34^, see Fig. 2). This procedure was performed on particles extracted with binning factors of 6 and 2 as well as unbinned (bin1), to determine the pixel size at which lamella thickness would potentially start to play a role. First, particles were extracted with a binning factor of 6 (corresponding to a pixel size of 7.3 Å), which reached the Nyquist-limit when more than 4000 particles were used, regardless of lamella thickness (see Fig. 2a). Thus, for lower magnification cryo-ET and STA, lamella thickness up to 255 nm does not negatively influence the resolution. Similar analysis with binning factor 2 and unbinned (corresponding to pixel sizes of 2.3 Å and 1.2 Å, respectively) clearly separated particles in the middle and thinner thickness groups that converged to higher resolution, from those extracted from thick lamellae (Fig. 2b, c). Interestingly, the particles from the middle-thickness group slightly outperformed those from the thinnest lamellae. We finally refined the unbinned (bin1) particles in M^35^, which is known to further improve the resolution by local refinement^10,17,19^. Indeed, we found a significant increase in resolution for each of the particle groups (see Fig. 2c, d). Both thin and middle thickness groups converged to the same resolution (4.5 Å), whereas the particles from the thicker lamellae converged to 4.9 Å. We conclude that lamellae with local thickness up to ca. 180 nm can be used without compromising the resolution. Thinner lamellae are not necessarily required for achieving the highest resolution.

**Fig. 2.**
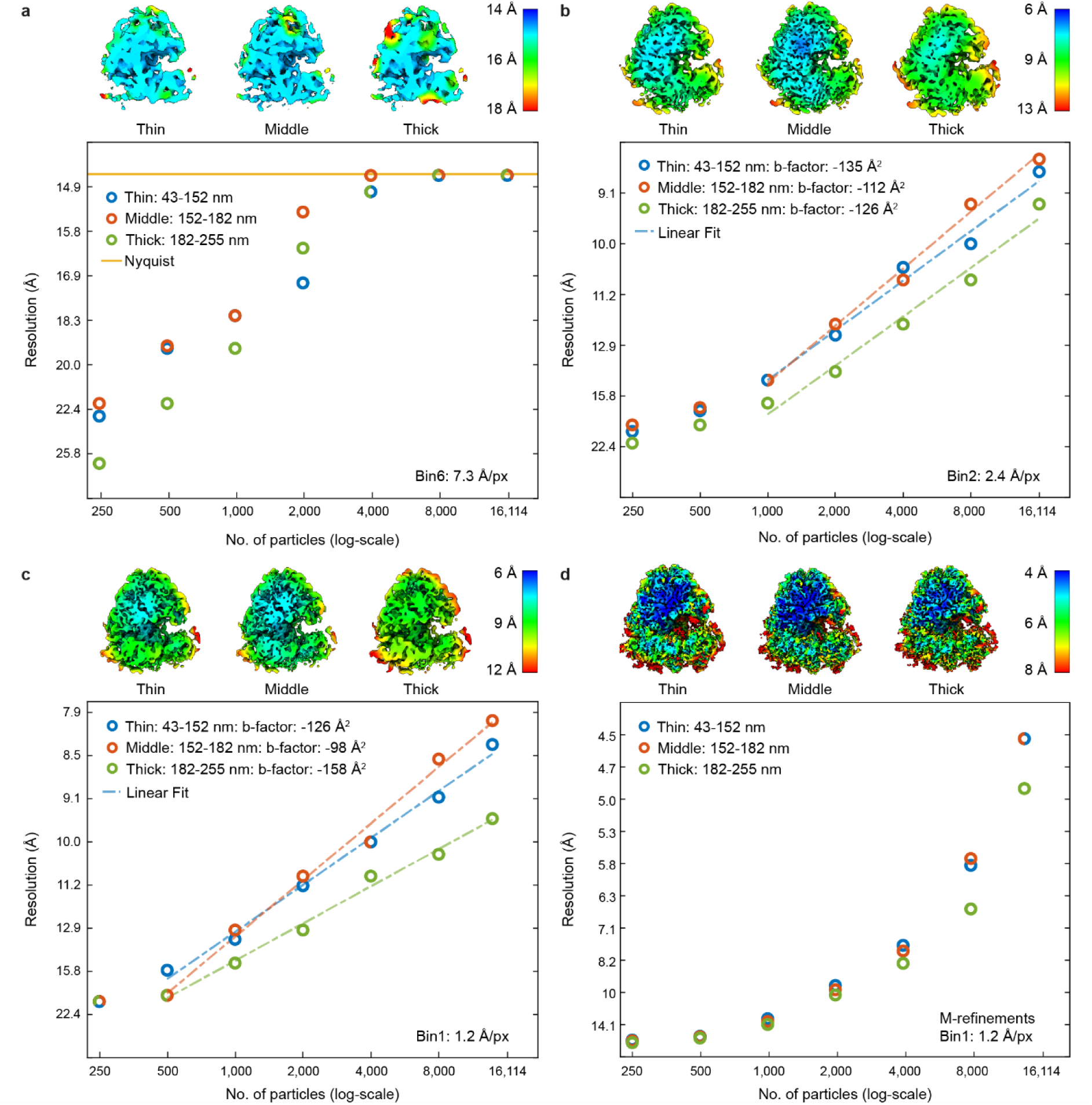
Local resolution density maps and resolution plots of different binning factors for lamellae with various local thickness. Resolution is plotted on the vertical axes (on square inverse scale). The number of particles on horizontal axes is on log-scale. Top: density maps of the largest particle subset for each thickness group. Bottom: resolution plotted against randomly selected subsets of particles. **a** The results for subtomograms extracted with binning factor 6. **b** Results for binning factor 2, with linear fit through the linear regime of the data^34^. **c** Results for unbinned data, processed with Relion 3D refinement only. **d** Results for unbinned data, further processed with M.

### Cryo-FIB milling causes structural damage to the sample up to ca. 30 nm from the surface

We then sought to quantify the depth and extent of structural damage caused by implanted ions and the collision cascade originating from cryo-FIB milling (see Fig. 3a). To control for the influence of thickness described above, we restricted the local lamella thickness to 140 – 190 nm for this analysis. To accurately extract particles at a given depth from the lamella surface, additional pre-processing steps that take into account the geometry of the sample were introduced (see “Methods” and Supplementary Fig. 2 for details). To quantify ion-damage in particles located close to the lamella surface, we selected particles from depths of 5 – 50 nm and compiled them in groups of 5 nm intervals, with 1000 particles in each group (see Fig. 3b). These were then extracted at binning factor of 2 (pixel size of 2.3 Å) and refined using Relion 3.1^32^. We found that particles centred around 5 nm from the surface averaged to 21.6 Å resolution. When particles were extracted from deeper within the lamellae, the resolution improved steadily until a plateau was reached at a depth of roughly 30 nm, where particles could be resolved up to 14.7 – 13.3 Å resolution (see Fig. 3d, e).

**Fig. 3.**
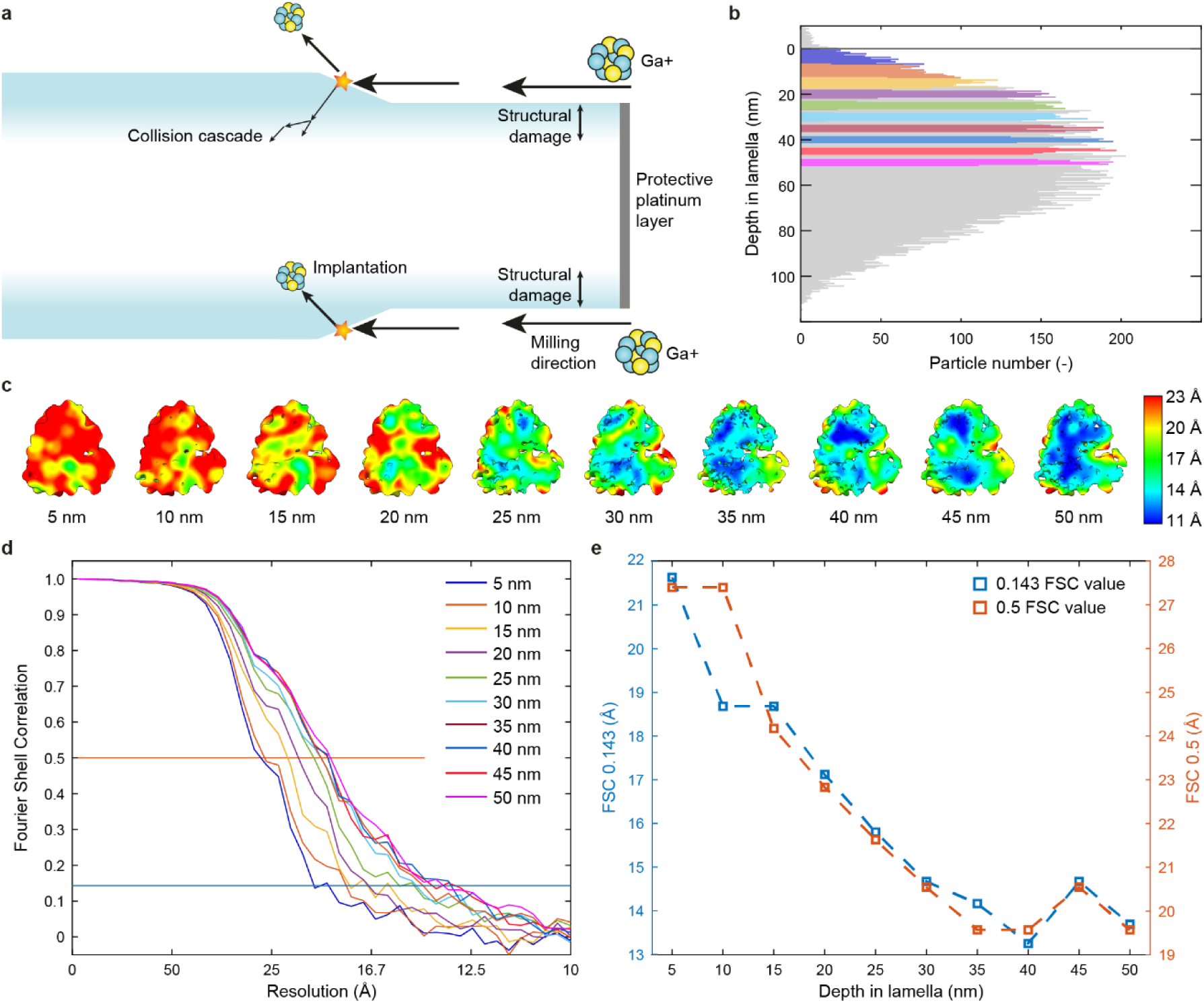
Ion-induced structural damage inside lamellae. **a** Milling geometry inside the cryo-FIB microscope. Upon milling and removing material, ions are implanted and cascade events cause structural damage in the lamella. **b** Histogram of all particles in the analysis, plotted with respect to the closest lamella surface (grey line). Colours indicate particles extracted for analyses for each of the depth-groups. **c** Subtomogram averages of 1000 particles from each depth, coloured according to local resolution. **d** Fourier shell correlation (FSC) of each depth group, using the colours from panel b. **e** Plot of the FSC value at cut-off values 0.5 and 0.143 for each particle subgroup.

These data clearly show that particles extracted closer to ca. 30 nm from either surface of the lamellae are of lower quality than those deeper inside the lamellae.

### Resolution from ribosomes slightly decreases from the front towards the back of the lamellae

As the cryo-FIB milling process is directed from the front towards the back of the lamellae, we analysed a potential quality difference of particles originating from specific regions. Again, to avoid any thickness effects, we limited the local thickness to 145 – 189 nm for this study. The data was split into particles from the damage layer (< 30 nm from the surface) and particles well away from damaged regions of the lamellae (> 60 nm from the surface). This led to 4 particle groups (front inner, front surface, back inner and back surface), of ca. 600 particles each, which were extracted and refined at binning factor 2 as before (see Fig. 4). While we detect a slight difference between particles from the front versus the back, both from inside (14.7 Å versus 15.8 Å respectively), as from the edge of the lamellae (18.7 Å versus 21.6 Å respectively), the quality of the data is dominated by the damage layer described above.

**Fig. 4.**
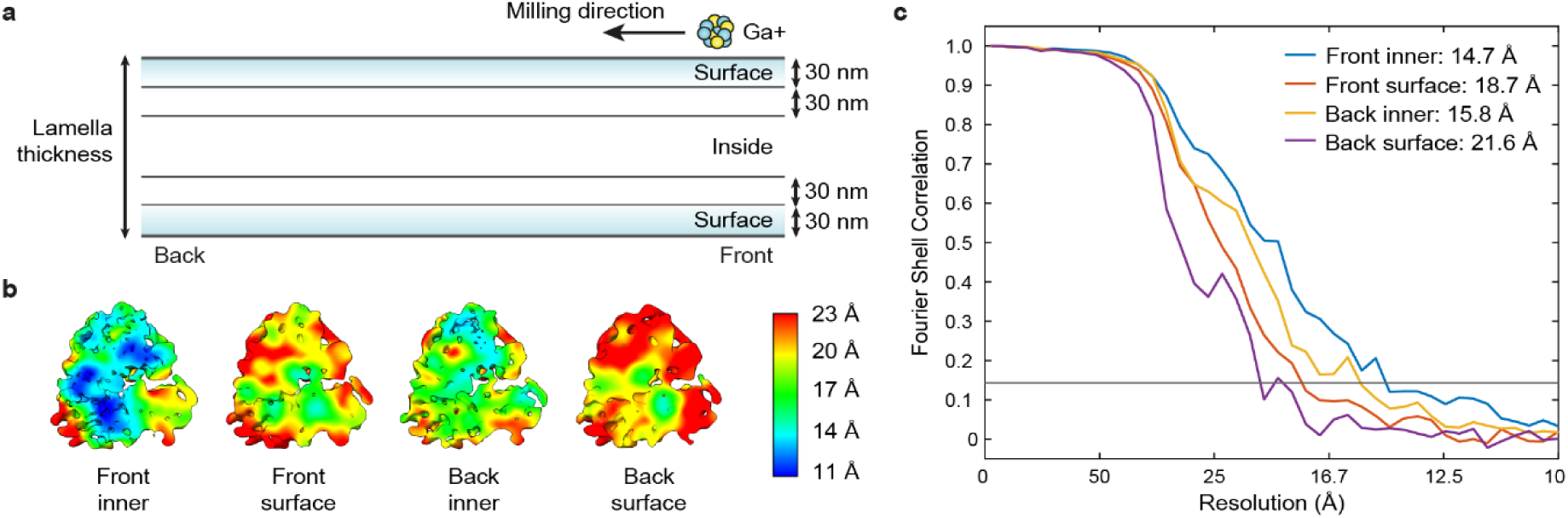
Resolution comparison between front and back of the lamella. **a** Milling geometry with respect to the front and back of the lamella. **b** Subtomogram averages of each of these particle groups, coloured according to local resolution. **c** FSC plot of the particle groups in b.

### High-quality particles average to higher resolution

Based on our findings, we extracted high-quality particles from the complete dataset from lamellae thinner than 180 nm and further than 30 nm away from the surface, which resulted in ca. 23,000 particles. To our surprise, these high-quality particles reached a similar resolution to an identical number of particles that were not filtered according to these quality criteria, 3.9 Å and 4.0 Å, respectively (see Fig. 5a). We hypothesized that although the randomly selected particles also contained particles of lower quality, this was most likely compensated by the relatively large total number of particles. Therefore, we also isolated two sets of ca. 5000 particles, high-quality and randomly selected from the whole dataset. Indeed, the high-quality particle set averaged to 6.1 Å resolution, while the randomly selected particles averaged to 6.9 Å (see Fig. 5b). Thus, using high-quality particles results in higher resolution. However, the reduced quality of particles close to the surface or from thicker lamellae can be compensated by increasing the number of particles.

**Fig. 5.**
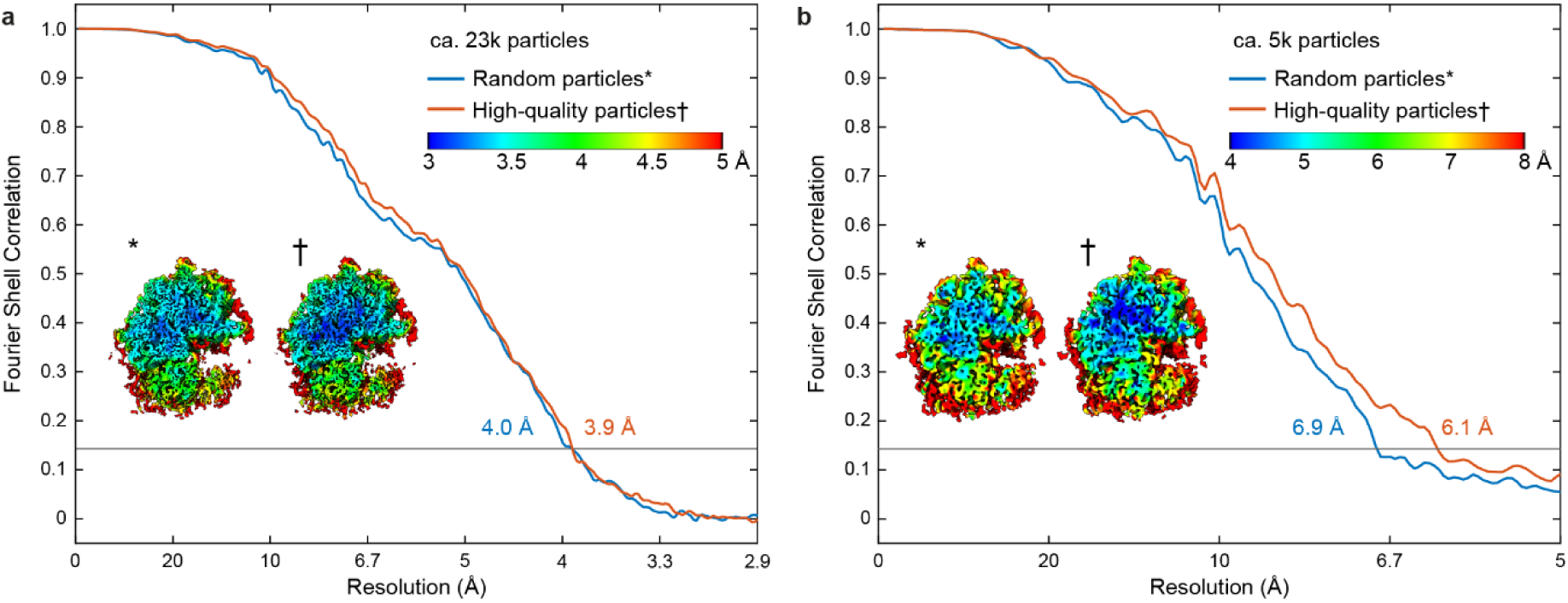
STA with high-quality particles compared to random particles. **a** FSC curve of 22,586 high-quality particles (red line) compared to 22,586 randomly selected particles (blue line). Inset: local resolution map of both sets. **b** FSC curve of 4795 high-quality particles (red line) compared to 4795 randomly selected particles (blue line). Inset: local resolution map of both sets.

## Discussion

Cryo-FIB milling of thick biological samples is gaining importance as *in situ* tomography of macromolecular complexes in (eukaryotic) cells becomes more widely used. Since its development for biological specimens and despite the recent surge in studies using this technique, no systematic assessment of its impact on structural output is available to date. In this work, we used ribosomes to benchmark the effect of cryo-FIB lamellae preparation on resolution attained by STA. Strikingly, we found that particles from thinner (up to 150 nm) lamellae did not reach higher resolution than from intermediate thickness (up to 180 nm). Previously, the effect of recent technological advances on the resolution of SPA cryo-EM has been investigated^9^, showing that 300 kV acceleration voltage, energy filters and super resolution detection strategies are particularly effective in thicker ice, all of which were exploited in the present study. Cryo-ET may benefit even more strongly from these technological advances than SPA, for example by enabling higher accuracy tilt-series alignment, thereby greatly reducing the thickness effects from the lamellae on resolution. Our finding that slightly thicker lamellae can be used without compromise on the attainable resolution is of great practical use, as the milling of very thin lamellae is not straightforward due to many technical challenges. Thicker lamellae can be comfortably used if high resolution is not the main focus.

In addition to the contribution of lamella thickness, we also assessed the effect of ion-damage during cryo-FIB milling on resolution and found that it causes structural damage to the sample up to ca. 30 nm from each surface. Particles from deeper layers in the lamella all averaged to roughly the same resolution. This result agrees well with the damage layer found when using argon ions using a plasma FIB^20^. We, however, note that the detected depth of 30 nm is comparable to the size of our probe, the ribosome, and thus may even be a conservative estimate. A recent study that used 2D template matching to probe the cross-correlation score between the sample and a high-resolution template reported a reduction of the signal-to-noise ratio for ribosomes up to ca. 60 nm from the lamella surface^25^. However, this assessment was not based on the achievable STA resolution and was performed on 2D images collected at high dose corresponding to SPA image acquisition parameters.

When a relatively small number of particles were averaged, high quality particles from lamellae with local thickness of less than 180 nm and located deeper than 30 nm in the lamella averaged to a significantly higher resolution. However, our dataset of ca. 23,000 particles as well as other previously published high-resolution datasets^17,19^ suggest that the thickness-effect and ion-damage are not a major concern for the cryo-FIB milling and cryo-ET workflow and can be compensated by including more particles in the average.

We note, however, that the ion-damage is only apparent when performing STA. Visually, all tomograms appeared undamaged, unlike for *e.g.* crystalline material^36^ and cryo-sectioning techniques^37^, so for morphological studies the ion-damage does not have to be considered.

Taken together, our systematic analysis shows the effect of cryo-FIB milling on the resolution achieved by STA. We found a minor loss of quality from particles originating from thicker parts of lamellae, or from close to the lamella surface, resulting in a decrease in resolution. However, for many cases, such as lower resolution averaging and morphological studies, these effects are negligible. Even for high resolution studies, we have shown that these effects can be countered by expanding the dataset to include more particles. These results show that cryo-FIB milling in general is not impeding any high-resolution structural studies.

## Methods

### Cryo-EM sample preparation

*D.discoideum* strain Ax2-214 cells were grown in HL5 medium (Formedium) containing 50 µg/mL ampicillin and 20 µg/mL geneticin G418 (Sigma Aldrich) at ca. 20 °C.

200 mesh Au grids with R1/4 SiO_2_ foil (Quantifoil) were glow-discharged for 90 sec at 0.38 mBar and 15 mA using a Pelco easiGlow device. Exponentially growing cells were diluted to a concentration of 3.3 x10^5^ cells/ml. A droplet of 100 µl cell suspension was placed on the glow discharged grids and cells were allowed to attach to the grid for 2-4 hrs at room temperature. Cells were subsequently vitrified by plunge freezing into liquid ethane using a Leica GP2 plunger. Lamellae were prepared by cryo-FIB milling with an Aquilos FIB-SEM (Thermo Scientific). Prior to milling, grids were coated with organometallic platinum layer using a gas injection system for 10 sec and additionally sputter coated with platinum at 1kV and 10 mA current for 20 sec. SEM imaging was performed with 10 kV and 13 pA current to guide the milling progress. Rough milling was performed using SerialFIB^38^ in three steps. Distance between milling patterns were reduced from 3 µm to 1.2 µm and 600 nm, using 500, 300 and 100 pA current for 300, 300 and 200 sec, respectively. Fine milling was performed manually at 30 pA, starting with the patterns 250 nm apart. The top pattern was kept at the same position, whilst the bottom pattern was raised in small steps to make the lamella thinner. All steps were carried out with 30 kV acceleration voltage. During polishing, intermittent SEM-scans were used to ensure the protective platinum layer was still intact. Lamellae were targeted to a thickness between 75 - 180 nm. Roughly half of the lamellae were sputter coated with platinum after polishing for 1 sec at 1 kV and 10 mA.

### Cryo-ET data acquisition

The Cryo-ET dataset was acquired in two microscope sessions, from 13 lamellae on two grids. Data were collected on a Titan Krios G4 microscope operated at 300 kV equipped with a cold FEG, Selectris X imaging filter, and Falcon 4 direct electron detector, operated in counting mode (all Thermo Scientific). Overview montages of individual lamellae were acquired with 3.0 nm pixel size to select suitable areas for tilt-series collection. Tilt-series were acquired using SerialEM (version 4.0.1) in low dose mode as 4k x 4k movies of 10 frames each and on-the-fly motion-corrected in SerialEM. The magnification for projection images of 105,000x corresponded to a pixel size of 1.223 Å. Tilt-series acquisition started from the lamella pretilt of +8° and a dose symmetric acquisition scheme^39^ with 2° increments grouped by 2 was used, resulting in 61 projections per tilt-series with a constant exposure time and targeted total dose of ca.120 e^-^/Å^2^. An energy slit with 10 eV width was inserted and the nominal defocus was varied between −2.5 to −4.5 µm. Dose rate on the detector was targeted to be ca. 6 e^-^/px/s at the untilted specimen on a representative position on a lamella.

### Image processing, template matching and classification

The motion corrected tilt-series were dose-filtered using Matlab-based scripts^40^ and cleaned by visual inspection. The dose-filtered tilt-series were then aligned through patch-tracking and reconstructed as back-projected tomograms with SIRT-like filtering of 10 iterations at a binned pixel size of 4.9 Å (both done in IMOD 4.11.5^28^). From the reconstructed tomograms 216 were selected by visual inspection. Tilt-series were rejected based on the tilt-images being very blurry, image acquisition was aborted for experimental reasons, or the reconstructed tomogram was extremely blurry. For compatibility with Relion^32^ and M^35^, the selected tilt-series were then reprocessed in Warp^30^ with the alignment obtained from IMOD (see Supplementary Fig. 1 for a complete overview).

For template matching, a ribosome average was generated using Relion from approximately 1000 manually picked ribosomes, with binning factor 10 (12.2 Å/px). Template matching was performed with the initial ribosome average on deconvolved tomograms with binning factor 10 reconstructed in Warp using STOPGAP^31^. For each tilt-series, the 800 highest cross-correlation peaks were selected, their coordinates exported and converted to a Warp-compatible star-file using the dynamo2m toolbox^41^.

The 168,000 positions determined through template matching were extracted as subtomograms in Warp at binning factor 6 (7.338 Å/px), and subjected to multiple rounds of 3D classification using Relion 3.1 to filter out junk particles. This yielded 48,342 ribosomal particles, which were used for successive processing steps, also see Supplementary Fig. 1.

### Particle curation and refinement for the thickness analysis

For the thickness analysis, the thickness of the lamellae was measured manually using IMOD. Three thickness-groups were created by equalizing the number of particles among these groups. This resulted in thickness ranges of: 43 – 152 nm, 152 – 182 nm and 182 – 255 nm, each containing 16,114 particles. Lists of particle indices referring to the particle starfile exported from STOPGAP, cross-referenced to the classification results, were created in Matlab, which served as the input for starparser^42^ to create separate star-files with particles for each thickness group. Separate Warp-projects were made for each of the thickness groups. To create Rosenthal-Henderson plots, subsets of particles (of 250, 500, 1,000, 2,000, 4,000, 8,000 and 16,114 particles) were created by randomly reducing the number of particles in the starfile in a sequential manner, using starparser^42^. The random selection of particles was checked to ensure unbiased thickness distribution in the subsets. For the unbinned data instead of 16,114 particles, only 13,840 particles could be extracted due to computational limits, however particle number between the thickness groups were kept identical.

Then subtomograms with binning factor of 6 (corresponding to a pixel size of 7.3 Å/px), 2 (2.4 Å/px) and unbinned subtomograms (1.2 Å/px) were extracted using Warp, and refined and post-processed in Relion 3.1. Refinement results from previous binning factors were used to re-extract subtomograms at lower binning to have better initial particle orientation and speed up further refinement runs. Next, based on the refinement of unbinned subtomograms in Relion, successive refinements were carried out in M (version 1.0.9). Multi-particle refinement of the tilt-series, geometric and CTF parameters were refined in a sequential manner until the resolution no longer improved, see Supplementary Table 1. Care was taken to first process the 250-particle subset, and then continue with the larger subsets in order to avoid influence of the multi-particle alignments from a refinement with higher particle numbers. Linear fits for the Rosenthal-Henderson plot were created in Matlab. Note that for the M-refined data, no linear regime was found to fit for a b-factor, due to M’s multi-particle refinement and because of the way the random selection of subsets were selected, the number of particles per tomogram increased, thus increasing the resolution in a non-linear manner.

### *In silico* straightening of ribosome coordinates for ion-damage analysis

To avoid thickness-effects only tomograms containing lamellae with local thickness in the range of 140 – 190 nm were selected. Firstly, the distance from each particle to the lamella edge had to be accurately determined. For various reasons, such as pre-tilt, offset between loading from the cryo-FIB microscope to the transmission microscope and inherent non-flatness of the lamellae, the cellular material is not aligned with the XY-plane in the reconstructed tomograms. Therefore, the ribosome coordinates were straightened within each tomogram, and the edges of the lamellae were determined. Using custom Matlab-scripts, a plane was fit through the raw ribosome template coordinates, prior to classification, see Supplementary Fig. 2a. For each raw ribosome coordinate, the Z-coordinate of the fitted plane was subtracted from the Z-coordinate from template matching, which resulted in a flattened lamella, roughly centred around Z = 0, see Supplementary Fig. 2b. From a histogram of all Z-coordinates from one tomogram, the position of both the top and bottom lamella edge were estimated as where the number of template matches drops off sharply around the edges of the lamella, see Supplementary Fig. 2b, c. Then, the distance from the ribosome coordinate to the closest lamella edge was calculated, where no distinction between top or bottom edge was made.

Particles originating from specific depths in the lamellae were selected as follows. 1000 particles most closely centred around 5 nm depth from the edge were first selected. This was then repeated with 5 nm intervals until 50 nm from the lamella edge, and for each depth 1000 closest particles were selected, using a custom Matlab-script and used as input for starparser. For the 5 and 10 nm distance group, there is an overlap in particles, as not enough particles were located so close to the edge of the lamella. Subtomograms were extracted with binning factor 2 using Warp, corresponding to a pixel size of 2.4 Å/px. These subtomograms were refined and post-processed with a solvent-mask in Relion for accurate Fourier shell correlation (FSC) comparison. Lastly, local resolution jobs were run in Relion.

For the analysis of the resolution between the front and back of the lamella, tomograms from the front and back were selected manually, chosen such that the thickness range was 145 – 189 nm for all. Then, particles were grouped into close to the surface (within 30 nm from the lamella surface), or particles that were > 60 nm away from the surface, to avoid any ion-damage. These 4 groups were processed like before, extracted with a binning factor of 2 using Warp and refinement, post-processing and local resolution jobs were run in Relion.

### High-quality particle selection and processing

To select high-quality particles from the entire dataset, first, tomograms with local thickness smaller than 180 nm were selected. Then, particles from the damage-zone (closer than 30 nm from the lamella surface) were excluded. These particles were extracted in Warp, refined in Relion, and successively refined in M, as described above. As comparison, tomograms and particles were randomly selected so that the number of particles were identical to the high-quality dataset, and these were processed in an identical fashion.

Then, for the smaller subset of high-quality particles, tomograms were randomly selected until 4795 high-quality particles could be extracted. These were compared with particles from randomly selected tomograms until 4795 particles could be selected. By selecting tomograms rather than particles, the effect of the multi-particle approach in M-refinement was maximised, and also mimicked realistic datasets. Again, particles were processed in Relion and M similar as before.

All density maps and local resolution maps were plotted using UCSF ChimeraX^43^.

## Data and materials availability

Relevant cryo-ET density maps generated in this study have been deposited in the EM Data Bank (EMDB) with the following access code: EMD-XXXXX.

All raw data has been deposited to the Electron Microscopy Public Image Archive (EMPIAR) database (accessing code EMPIAR-XXXXX), including tilt-series alignment data, pre-processing data, raw template matching locations and particle locations.

### Acknowledgements

This project has been made possible in part by grant number 2021-234666 from the Chan Zuckerberg Initiative DAF, an advised fund of Silicon Valley Community Foundation and by the Max Planck Society. We thank Mark Linder from the Central Electron Microscopy facility of the Max Planck Institute of Biophysics for technical support. We thank Patrick C. Hoffmann, Huaipeng Xing, Sergio Cruz-León and Tomáš Majtner for useful discussions. We thank Özkan Yildiz, Juan F.Castillo Hernandez, Andre Schwarz, Erin Schuman and the Max Planck Computing and Data Facility for support with scientific computing. We thank Stefanie Böhm and Patrick C. Hoffmann for critically reading of the manuscript. All data in this manuscript was acquired at the Central Electron Microscopy Facility at Max Planck Institute of Biophysics.

## Author contributions

MWT, GH, MB and BT conceptualised the project and developed the methodology. MWT and SW acquired the data. MWT and JPK processed and analysed the data. MWT, MB and BT wrote the manuscript, and all co-authors edited the manuscript. GH, MB and BT supervised the project.

## Supplementary Figures

**Supplementary Fig. 1.**
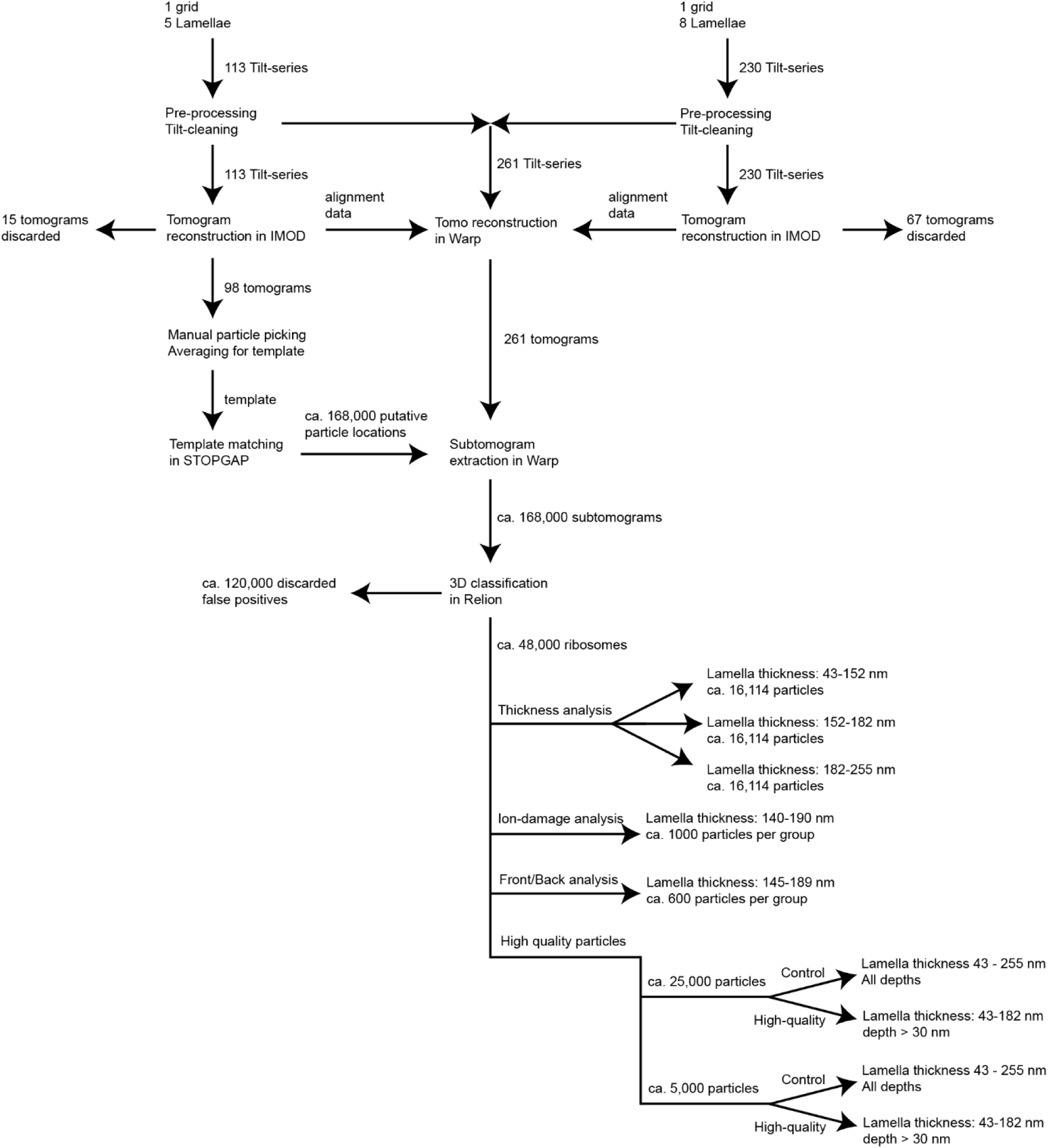
Processing workflow.

**Supplementary Fig. 2.**
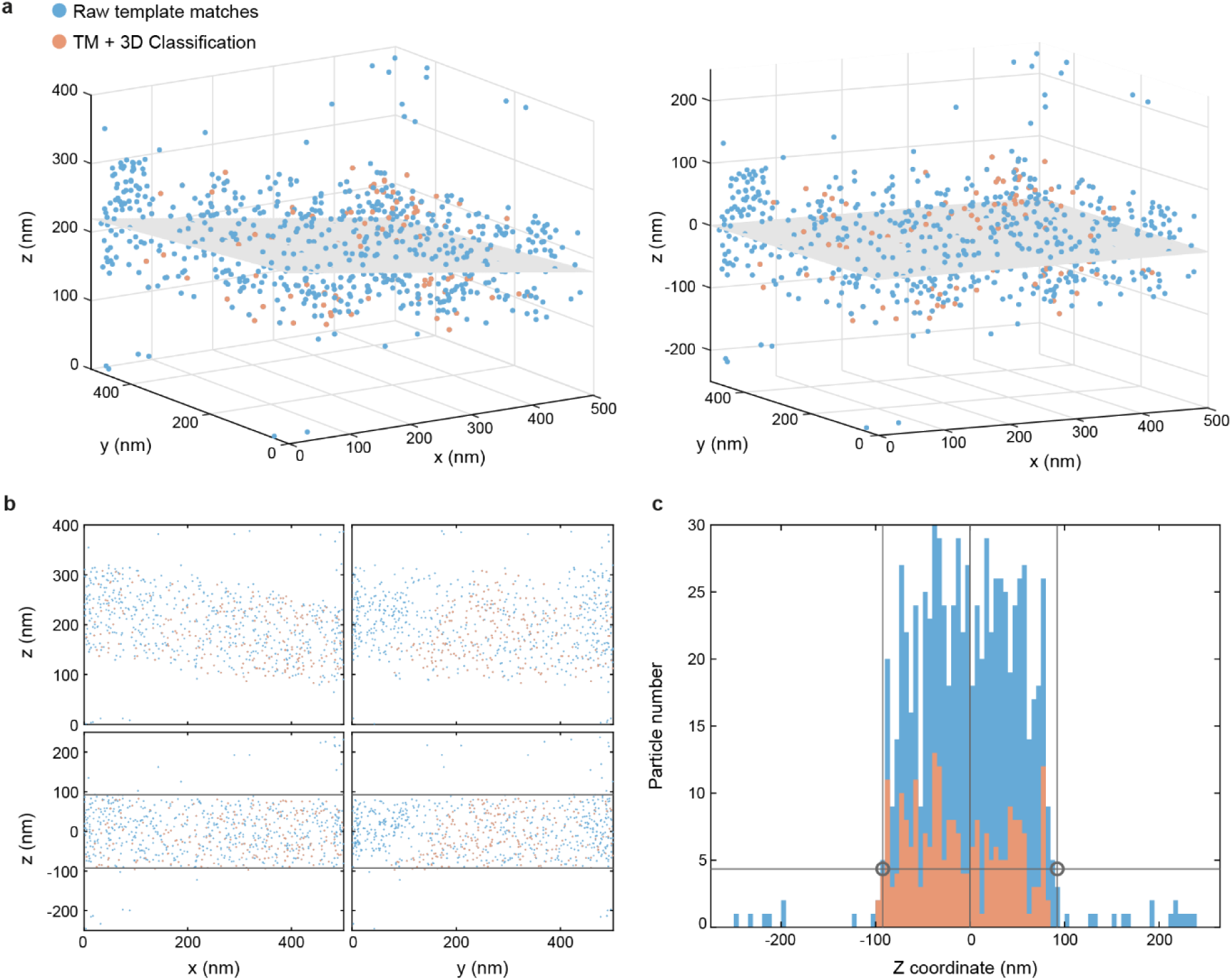
*In silico* straightening of lamellae and ribosome coordinates to find lamellae edges. **a** Locations of ribosome template matches from a single tomogram are used to fit a 2D plane (left) which is used to straighten the lamella (right). **b** X-(left) and Y-(right) ribosome coordinates before (top) and after (bottom) straightening. The grey lines represent the automatically detected edges of the lamella. **c** Histogram of straightened z-coordinates of the raw (blue) and classified (red) template matches. Lamella edges were determined based on the drop-off value of the coordinates, indicated with black circles.

**Supplementary Table 1.** M refinement iterations. Iteration step 3 was only performed when the resolution was at least 7.0 Å. Iteration steps 3-5 were only performed when the resolution still improved after performing the previous iteration step

| Iteration step | Geometry |  |  |  |  | CTF |  |
| --- | --- | --- | --- | --- | --- | --- | --- |
|  | Image warp grid | Particle poses | Doming | Stage angles | Volume warp grid | Defocus | Grid search |
| 1 | 3 × 3 | ✓ | - | ✓ | 3 × 3 × 2 × 10 | - | - |
| 2 | 6 × 6 | ✓ | ✓ | ✓ | 6 × 6 × 8 × 10 | - | - |
| 3 | - | - | - | - | - | ✓ | ✓ |
| 4 | 7 × 7 | ✓ | ✓ | ✓ | 7 × 7 × 8 × 10 | ✓ | ✓ |
| 5 | 10 × 10 | ✓ | ✓ | ✓ | 10 × 10 × 12 × 10 | ✓ | ✓ |

